# A scale-dependent measure of system dimensionality

**DOI:** 10.1101/2020.12.19.423618

**Authors:** Stefano Recanatesi, Serena Bradde, Vijay Balasubramanian, Nicholas A. Steinmetz, Eric Shea-Brown

**Author notes:** These authors contributed equally. These authors share senior authorship (Vijay Balasubramanian, Nicholas A. Steinmetz, Eric Shea-Brown).

## Abstract

A fundamental problem in science is uncovering the effective number of dynamical degrees of freedom in a complex system, a quantity that depends on the spatio-temporal scale at which the system is observed. Here, we propose a scale-dependent generalization of a classic enumeration of latent variables, the Participation Ratio. We show how this measure relates to conventional quantities such as the Correlation dimension and Principal Component Analysis, and demonstrate its properties in dynamical systems such as the Lorentz attractor. We apply the method to neural population recordings in multiple brain areas and brain states, and demonstrate fundamental differences in the effective dimensionality of neural activity in behaviorally engaged states versus spontaneous activity. Our method applies broadly to multi-variate data across fields of science.

---

In many branches of science, complex systems are characterized by the simultaneous values of a large number of observables evolving over time. For example, in cell biology, the operational state of a cell may be summarized by the expression levels of myriad proteins. Likewise, in neuroscience, the instantaneous activity levels of the many neurons in a brain region summarize the state of the system [1]. The dynamics of these systems can be much lower dimensional. For example, at the coarsest scale, the overall dynamics of a brain area may just be described by a slow fluctuation in the mean neural firing rate, i.e., by a single dynamical variable [2, 3]. At some intermediate scale, the same dynamics could consist of a certain number of characteristic firing patterns evolving smoothly on a fixed *d*-dimensional curved manifold embedded in the state space. If we knew the underlying dynamical system, we could derive from first principles the relevant manifold at each scale of dynamics. But in many of the most exciting complex systems that are becoming accessible to experimental study, our goal is to *discover* the dynamical system, a task that starts by determining the necessary number of effective latent variables, i.e., the dimensionality of the system.

One approach to this problem has its roots in basic point-set topology. In this approach, we recognize that a manifold is *d*-dimensional if we find that the number of uniformly sampled points within a region of characteristic length *L* scales as *L*^*d*^. This fact leads to definition of the *capacity dimension D*_0_ [4] in terms of the number *n* of Euclidean boxes needed to cover the system’s trajectory in its embedding space: 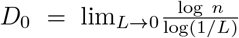. This intuitive notion of dimension is difficult to compute in more than three dimensions [5], and faces the challenge that sampling of dynamical systems is often too coarse to directly estimate the *L* → 0 limit [6–8]. A variation on this idea that is easier to estimate from high dimensional data is the *Correlation dimension D*_*corr*_, [9] which is determined by the scaling of the number of pairs of data points with separations less than *r*, in the *r* → 0 limit. In this way, the capacity dimension and the Correlation dimension both give local, fine-scale measures of dimension [9, 10].

A second class of approaches to the problem of measuring dimensionality starts with the correlation matrix between observations. There are many techniques related to the classic Principal Components Analysis, in which the effective dimension is defined as the number of eigenmodes of the correlation matrix of the data that are sufficient to capture most of the variance. A substantial literature analyzes how to choose a threshold that determines when “most” of the variance has been captured [4, 11], but this choice leads to a certain arbitrariness, and makes it challenging to define the dimension associated to different scales of observation [12, 13]. A more natural way to define a dimension from the correlation matrix is by computing the Participation Ratio [1, 14] of the eigenvalues, namely, a ratio of the square of the first moment and the second moment of the probability density function of the eigenvalues [15]. As we will discuss, this quantifies the number of effective dimensions along which the data are spread. In what follows, we will generalize the Participation Ratio dimension to measure the effective dimension at different scales of observation, and will show that this quantity interpolates between the Correlation dimension at small scales and the Participation Ratio dimension globally. We will further show that the quantity has an intuitive meaning when applied to known dynamical systems, and then use it to extract insights into the structure of neural population activity in different brain areas and brain states.

Consider a set of *T* observations of *N* observables ***x***(*i* = 1..*T*) sampled from an underlying probability distribution *ℳ*. In a dynamical systems context, ***x***(*i*) could be the *i*^th^ point generated by an iterative process ***x***(*t* + *τ*) = *ℱ*(***x***(*t*)) with fixed or varying initial conditions. But, more generally, is the distribution of the data regardless of how it was generated. The empirical covariance matrix ∑ over the observations ***x***(*i*) is 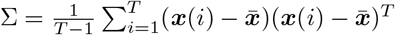 a *N × N* matrix, where the average 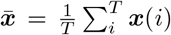. The eigenvalues of the covariance matrix ∑ are *λ*_*i*=1..*N*_ and the associated spectral density in the continuous limit is *ρ*(*λ*). The Participation Ratio dimension, *D*_*P R*_, is defined as the ratio between the second and the first moment of the spectral density [15]:

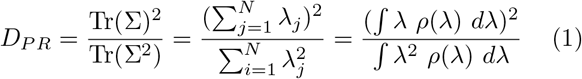

This object is a measure of concentration of the eigenvalue distribution and quantifies how many eigenmodes are necessary to significantly capture the overall distribution of observations *ℳ*, similarly to using the number of eigenmodes of ∑ that are sufficient to capture most of the variance.

To extend *D*_*PR*_ to a measure of dimensionality at different scales, we first compute, for each point ***x***_*i*_, the local covariance matrix of points up to a distance, *r*. In local Principal Components Analysis (lPCA), the dominant eigenvectors of this matrix determine the local subspace in which the distribution ℳ is localized. Computing the Participation Ratio dimension of this local covariance gives a measure of local dimension. Averaging this measure over all starting points yields the averaged quantity *D*_*P R*_(*r*) which is an effective dimension of the manifold up to the scale *r*.

We can compare this scale-dependent notion of effective dimension to another notion that arises from a generalization of the Correlation dimension that is defined as follows [16, 17]. Let *d*_*ij*_ = ‖***x***_*i*_ - ***x***_*j*_‖ be the distance between any two sampled observations, with the correlation integral at distance *r* defined as 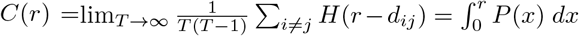, where *H* is the Heaviside step function and *P* (*x*) is the distribution of pairwise distances. We expect that *C*(*r*) ∝ *r*^*d*^, at least for small values of *r*, where *d* is the dimension of the manifold which supports the data distribution *ℳ*. Thus we defined 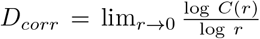. Although *D*_*corr*_ is defined in the *r* → 0 limit, this quantity is sometimes extended for a general *r* as the log-log derivative of the correlation

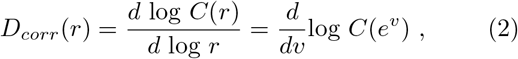

where *v* = log *r* [4].

In practice, for empirical observations where the number of measurements *T* is finite, a robust estimate of the Correlation dimension requires a test of covergence. When the dimension is relatively stable across a range of scales, one looks for a plateau in *D*_corr_(*r*) [16]. This definition of dimension has been very successful in treating strange attractors in chaotic systems in the sense that the Correlation dimension (i.e., lim_*r→*0_ *D*_corr_(*r*)) agrees with the well-known Lyapunov dimension which controls dynamical properties of the system [18, 19]. If we apply the generalization (Eq. (2)) across scales *r >* 0 we encounter a sampling challenge when the manifold is high dimensional. Because of this, even if the manifold has a fixed dimension across scales, it can be difficult to find the plateau in *D*_corr_(*r*) that is a sign of robust estimation [6, 8, 20]. The problem is compounded if we expect the effective dimension to vary with scale in the first place because we would then need enough data to compute a bootstrap measure of the reliability of the scale-dependent estimate of dimensionality.

We can get an intuitive understanding of the Correlation dimension *D*_*corr*_(*r*) and of the Participation Ratio *D*_*P R*_(*r*) by evaluating them on two known examples, the Lorentz attractor and the 2d Spiral with added local 2d noise (Fig. 1) [4]. In the case of the Lorentz attractor, both *D*_*P R*_(*r*) and *D*_*corr*_(*r*) recover an effective dimension just bigger than 2 locally (i.e. as *r* → 0), matching expectations from dynamical systems analysis [19]. However, as the scale, i.e. *r*, increases, we see that the two measures have very different behavior. At large scales the Correlation dimension declines to 0. This is because the Lorentz attractor is a compact manifold and if we examine it naively at very large scales it looks pointlike. At a technical level, this follows from the derivative with respect to the scale *r* of the correlation integral in the definition of the Correlation dimension (Eq. (2)). By contrast the participation ratio dimension remains roughly constant across scales. This is because this quantity includes correlations between every pair of datapoints up to the scale *r*, and not just at this scale, and then includes an adaptive rescaling of lengths (the denominator of the definition in (Eq. (1)), so that the effective dimension remains comparable across scales. We see a similar pattern in the analysis of spiral with local noise in Figs. 1d to 1f, where again the Correlation dimension declines to zero at large scales while the Partiticipation Ratio dimension correctly captures the essentially two-dimensional structure. The effective rescaling that leads to a finite Participation Ratio dimension even at large scales is especially important for analyzing high dimensional, finite size datasets. In this case, the data is always sub-sampled simply because it is difficult to explore a large number of dimensions, and thus it is hard to be certain whether the Correlation dimension is small because the data manifold is really low dimensional, or because of the asymptotic effect of *D*_*corr*_ going to zero at large *r*. Thus the Participation Ratio dimension is likely to give more useful information.

**FIG 1.**
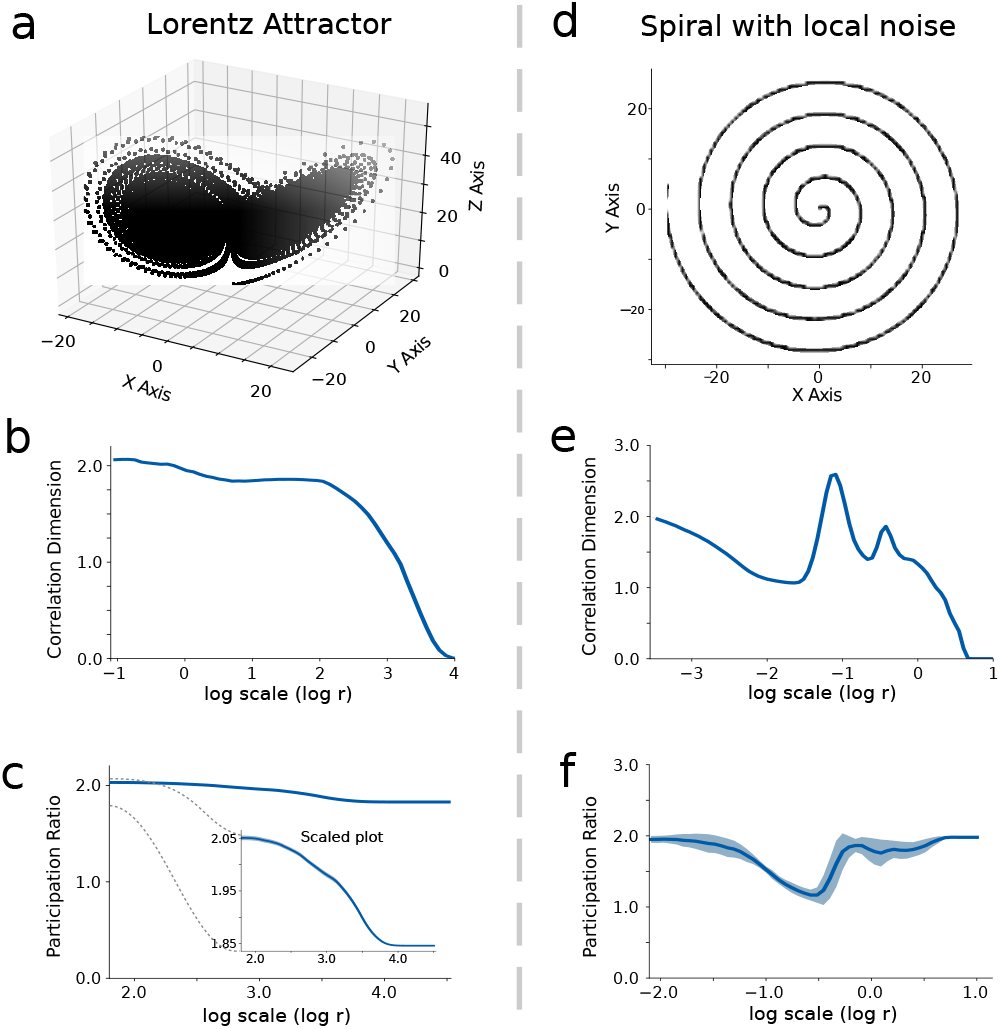
Correlation and Participation Ratio dimensions on benchmark examples. a) Distribution of 10^5^ points sampled from the Lorentz attractor. b) Scale-dependent Correlation dimension of the Lorenz attractor. In the small scales limit, the Correlation dimension matches the expectation from dynamical system analysis, the Lyapunov exponent 2.05, while in the opposite limit, it goes to zero. c) Scale-dependent Participation Ratio dimension of the Lorentz attractor. The dimension at every scale is averaged over 100 randomly sampled points. Error bars represent the standard error over such statistics. Inset: zoomed in scaled version of the same plot (scaled y-axis, same x-axis). This dimension is a good estimator of the dimension for several orders of magnitude. e) Distribution of 10^5^ points sampled from the the 2d spiral with local 2d noise. This dataset is locally two dimensional at small scales due to the added noise, then one dimensional at intermediate scales and finally two dimensional at larger scales. e) Correlation dimension of the 2d spiral wave interpolates between 2 to 1 at small scale but then decays to zero non-monotonously for larger scale. f) Participation Ratio dimension of the 2d spiral. This quantity provides a good predictor of the real dimensionality of the system on the full range of scales that we studied.

To understand the relation between *D*_*P R*_ and *D*_*corr*_, it is helpful to consider these quantities at small data separations *r*, in which case we take a tangent space approximation to the data manifold. The data themselves will be distributed on this tangent space (or narrowly around it if there is noise). Additionally, both *D*_*P R*_ and *D*_*corr*_ are constructed from the second moments of data. Thus, locally, we can consider a second order moment approximation of the data

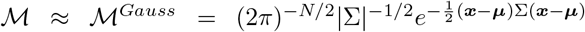

where ***µ*** is the mean of the distribution and ∑ the covariance matrix. Let us consider the rotated and centered coordinate system where ***µ*** ≡ 0 and ∑ is diagonal. In this presentation, sampled vectors ***x***_*i*_ have coordinates 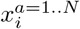 that are normally distributed *x*^*a*^ ∼ *N* (0, *λ*_*a*_) where *λ*_*a*_ is the eigenvalue of the covariance ∑ corresponding to axis *a*. Thus the Participation Ratio dimension is 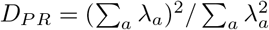.

To compare this to the Correlation dimension we want to compute the distribution *P* (*x*^2^) of squared Euclidean distances 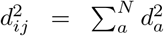 between sampled points ***x***_*i*_ and ***x***_*j*_. Note first that the difference squared 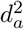 along the a^th^ axis of vectors ***x***_*i*_, ***x***_*j*_ sampled from *ℳ* is distributed according to a Gamma distribution: 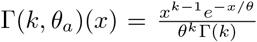 with *x >* 0 where *k* = 1*/*2 and *θ*_*a*_ = 2*λ*_*a*_. The overall distribution of Euclidean squared distances, *P* (*x*^2^), is then the convolution of independent Gamma distributions. It is approximately [21, 22] a Gamma function with parameters (*k*_*s*_, *θ*_*s*_) given by the Welch-Satterthwaite equations [23, 24] which simplify in our case to:

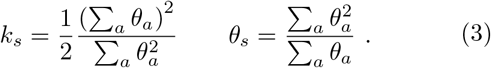

The correlation integral is then 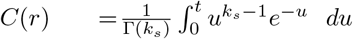 where *t* = *r*^2^*/θ*. Then, for *r* ≪ *θ*_*s*_, the exponential is approximately 1, and we obtain that 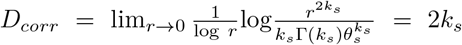. This yields:

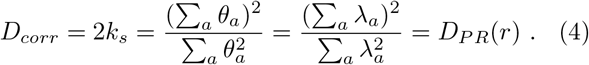

We conclude that the Participation Ratio *D*_*P R*_ coincides with the system’s Correlation dimension *D*_*corr*_ at small scales. In this sense *D*_*P R*_(*r*) agrees with a well-accepted local definition of dimension and generalizes this notion across scales in a different manner than the extension *D*_*corr*_(*r*) in (Eq. (2)).

To illustrate the relationship between the two studied measures we numerically computed *D*_*corr*_(*r*) and *D*_*P R*_(*r*) as a function of the scale *r* for isotropic multidimensional Gaussians, validating the analytical results (Fig. 2a and Fig. S1). While *D*_*corr*_ decreases at larger scales, *D*_*P R*_ remains constant. Furthermore we see how, limiting the sampled statistics to *N* = 50000 points it is not possible to achieve a plateau in *D*_*corr*_, even for a dimensionality of 10 – so that evaluating the dimensionality of the system from the plot of the Correlation dimension in Fig. 2a(left) is difficult if not impossible. In the case of a two dimensional Gaussian with increasing elongation along one of the two coordinate axes, Fig. 2b, the Correlation dimension captures well the local dimension of 2 for *r →* 0 but doesn’t allow one to quantify the increasing skewness of the distribution. Similarly *D*_*P R*_ allows the quantification of both its local dimension of 2 and its skewness, which induces a lower dimension at larger scales. A third important example, Fig. 2c, is the case of a scale-free system where the distribution of eigenvalues of the correlation matrix is power-law.

**FIG 2.**
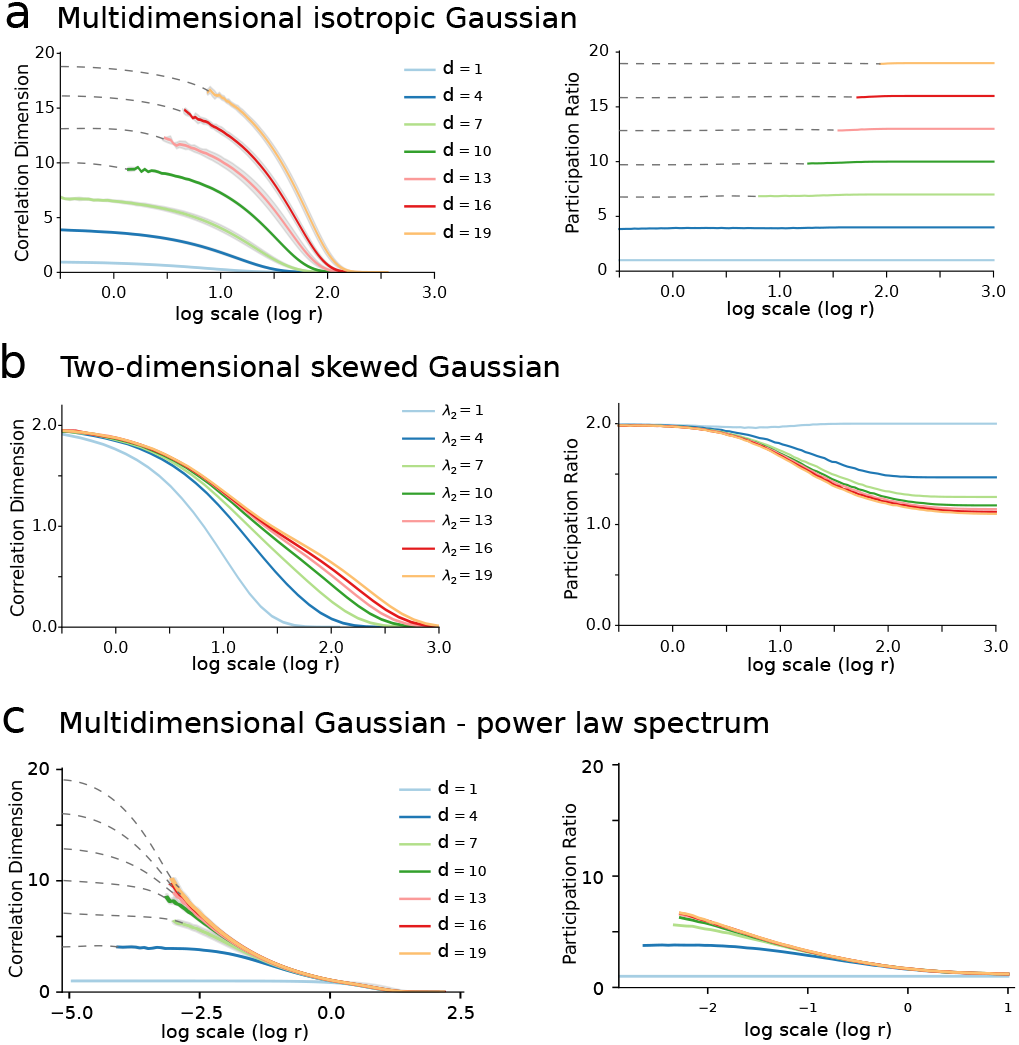
a) (Left) scale-dependent Correlation dimension and (Right) scale-dependent Participation Ratio of the multidimensional isotropic Gaussian distribution. Different lines correspond to increasing dimension *d*, cf. legend. b) (Left) scale-dependent Correlation dimension and (Right) scale-dependent Participation Ratio for the two-dimensional skewed gaussian distribution. The first eigenvalue of the diagonalized covariance matrix ∑ of this distribution is *λ*_1_ = 1 while the second varies according to the legend determining the elongation of the distribution. c) (Left) scale-dependent Correlation dimension and (Right) scale-dependent Participation Ratio of the multidimensional Gaussian distribution with scale-free power law spectrum. Different lines correspond to increasing dimension *d*, cf. legend. In this last case, the dimensions have eigenvalues which are distributed according to a power-law with *α* = −4 so that the k-th eigenvalue is *λ*_*k*_ = *k*^*−α*^. For each case under study, 50000 points were randomly sampled.

Systems with such properties are often deemed to be near criticality and are studied in a number of contexts. In this case *D*_*P R*_ is more effective than *D*_*corr*_ as it saturates at a finite value that depends only on the power law coefficient of the scale-free spectrum, while the Correlation dimension doesn’t display any plateau. Plugging a scale-free spectrum, *λ*_*a*_ = *βa*^*−α*^, into the formula for *D*_*P R*_ we find:

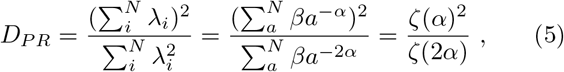

where *ζ*(*α*) denotes the Riemann Zeta function evaluated in *α*. For these systems *D*_*P R*_ and −*α* are in one to one correspondence, Fig. S2. The higher is the value of *α*, the smaller is the dimensionality *D*_*P R*_ of the system [25].

We applied our method of quantifying scale-dependent system dimensionality to the neural activity of thousands of simultaneously recorded neurons [26]. Massive neural recordings of this kind are becoming possible because of technological advances [27, 28], and a pressing question is how to uncover and characterize the dimensionality of the underlying neural dynamics [1, 25, 29]. However, it has not yet been determined whether the relevant dimensionality differs across scales. Here, we illustrate how the scale-dependent Participation Ratio dimension *D*_*P R*_(*r*) reveals the differences in dimensionality across scales, brain areas, and brain states.

To this end, we analyzed multielectrode Neuropixels recordings in four brain regions: visual thalamus, visual cortex, frontal cortex and midbrain. The first two are primarily sensory areas while the second two are involved in decision making [26, 30, 31]. We compared across two conditions: (i) spontaneous, where the animal was awake but with no specific task, (ii) engaged, where the animal performed a two-alternative forced choice task (cf. Suppl.Mat.) [26]. For each of the 37 recording sessions we analyzed the activity of all neurons, subdividing them in groups of 100 neurons each to allow comparison across recording sessions which sampled variable numbers of neurons. For each region and group of neurons we computed vectors of neural action potential counts in 100 ms bins. We then analyzed *D*_*corr*_(*r*) and *D*_*P R*_(*r*) with varying scale parameter *r* (Fig. 3a).

**FIG 3.**
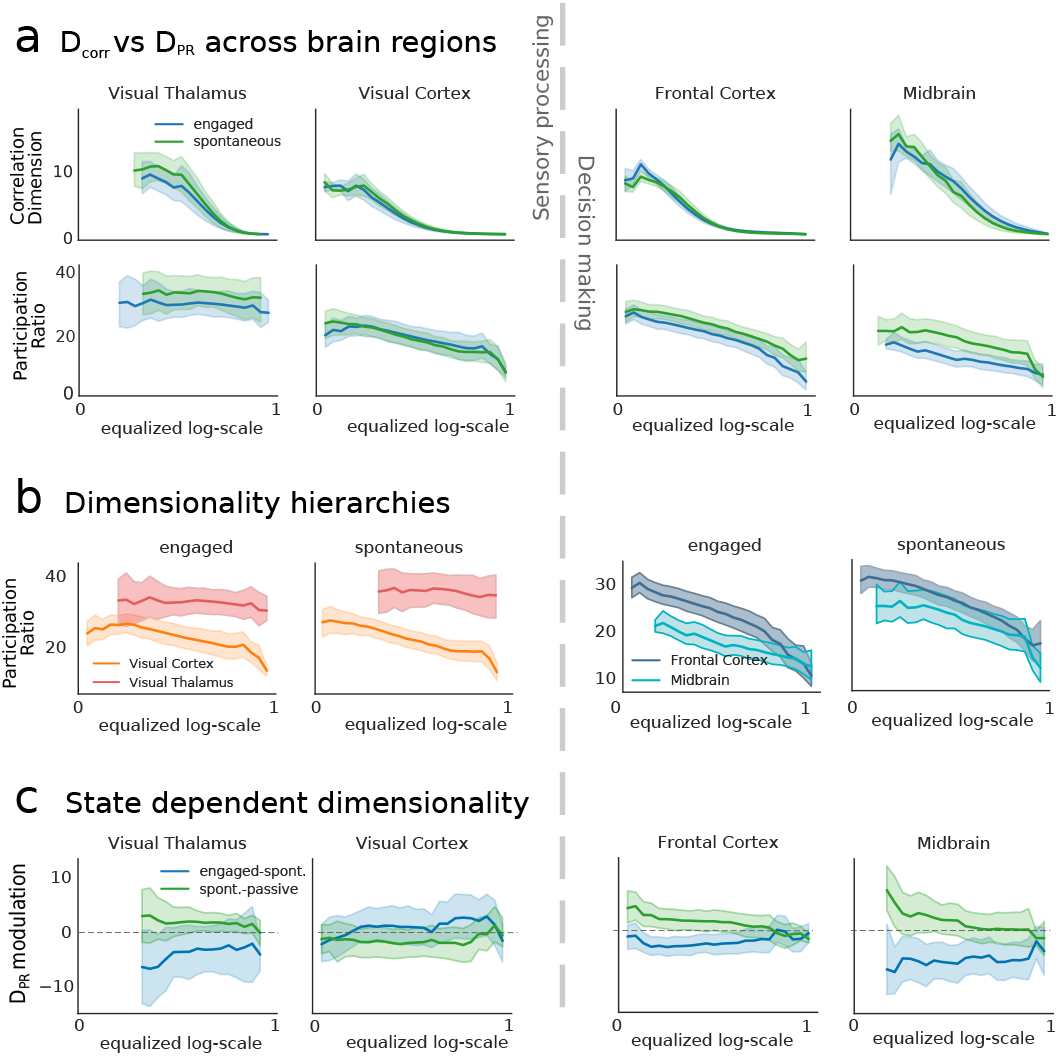
State dependent dimensionality of neural activity. a) Scale-dependent dimensionality analysis across brain regions. For each brain region the scale dependent Correlation dimension and Participation Ratio has been computed and averaged across sessions (cf. Suppl.Mat.). The scale varies from local scale (value of 0) to global scale (value of 1). Each line represents a separate condition as reported in the legend. Shaded areas represent 95% confidence interval across experimental sessions. b) Dimensionality for different brain regions. Error bars represent 95% confidence interval across experimental sessions. c) Scale-dependent dimensionality modulation Δ^*AB*^ between conditions A and B across brain regions. Shaded areas represent 95% confidence interval. The blue line represents Δ^*AB*^ with A equals to engaged and B equals to spontaneous, while the green line represents the conditions A, spontaneous, and B, passive.

As expected from the analysis above, *D*_*corr*_(*r*) rapidly converged to 0 when increasing the scale *r*. However, *D*_*P R*_ was more stable, enabling us to analyze dimensionality across a wide range of scales. Intriguingly, this dimensionality depended systematically on scale in visual cortex, midbrain, and frontal cortex – decreasing at larger scales (*D*_*P R*_(*r*_*min*_)*/D*_*P R*_(*r*_*max*_) = 1.63 ± 0.05 mean ± s.e.m., p=2.9 · 10^*−*4^, t-test) – but was roughly constant across scales in the visual thalamus (Fig. 3a). We emphasize that *r* is computed in terms of distances between neural response vectors, and thus represents a scale in the functional space of neural activity, rather than physically on the cortical sheet. Thus, we see that there are more degrees of freedom separating nearby vs distant neural responses in many areas, but that tendency this is not universal in the brain – in the visual thalamus, responses were instead equally rich across small and large scales.

Comparing dimensionality across brain regions in the visual stream (Fig. 3b), we observed that at every scale we studied the thalamus had a significantly higher dimensionality than visual cortex (p=2.7 · 10^*−*60^, t-test). Note that while anatomical expansion presumably entails a higher dimensionality of image representation in visual cortex than in thalamus [32], here we measured not the dimensionality of image representations, but instead the dimensionality of ongoing activity patterns during conditions with limited visual stimuli. We also found that frontal cortex had a higher dimensionality than midbrain in the engaged, but not spontaneous condition – and this difference was prominent at smaller scales only accessible by means of our analysis.

To directly compare changes in neural dimensionality between behavioral conditions, we analyzed the relative dimensionality difference, defined as 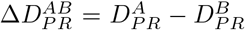, between conditions A (engaged) and B (spontaneous) (Fig. 3c, blue line). This difference was significantly different than zero in both frontal cortex and midbrain (p=2.9 · 10^*−*26^ and p=7.0 · 10^*−*27^ respectively, t-test). Specifically, task-driven neural activity was lower dimensional than spontaneous, and this difference was more prominent at small and intermediate scales. The same significant difference was not found in visual areas. To further evaluate the dependence of dimensionality on behavioral task, we compared against a third condition, called “passive”. In this condition, the same stimuli previously presented during the decision task were presented in randomized order without any behavioral response from the subject. The difference 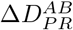 between spontaneous and passive was not significantly different from zero (green line). This highlighting the stronger dimensionality modulation induced by task-evoked activity, across a range of scales.

We concluded that our method of quantifying scale-dependent system dimensionality can extract new and fundamental features of neural activity patterns recorded across the brain. Specifically, dimensionality is not always characterized with a single number, but can vary systematically with scale across the brain: while some areas had the same dimensionality across all scales (Thalamus) others (cortex and midbrain) displayed a decrease at large scales. Nevertheless, clear hierarchies in dimensionality across brain areas exist, and hold across wide ranges of these scales. Finally, we showed how and where dimensionality varies with state.

Our work establishes the Participation Ratio as a theoretically grounded measure of dimensionality for complex systems at all scales. This has the potential to impact analyses in the many fields of science that have independently arrived at the Participation Ratio as a measure of ‘effective’ instrinsic dimension. In physics this ratio was first introduced in atomic spectroscopy [33, 34] and then used as a measure of localization in condensed matter physics [15]. In quantum information the same quantity is called the “purity”, and measures the degree of mixedness of quantum states. In economics and sociology a similar quantity, the Herfindahl–Hirschman Index, measures the market concentration of an industrial sector [35, 36]. In sociology the related Simpson index quantifies diversity [37], while in politics it is a measure of the effective number of parties [38]. In machine learning the same quantity serves as a metric of expressivity for learning kernels [39], and in neuroscience, this quantity is used to measure the dimensionality of neural activity [1, 40, 41]. The ubiquitous use of the Participation Ratio to measure the effective dimension of a complex dynamical system underscores its efficacy. However, as we have emphasized, complex systems have different behavior if observed at different scales and so their intrinsic dimension at different scales need not be a constant. The scale-dependent Participation Ratio as a measure of “running dimension” capturing the effective number of latent degrees of freedom required to summarize the observables at different scales. Our method is amenable to further theoretical analysis as *D*_*P R*_ is a simple functional of the second order statistics of a system can be analytically computed in many interesting theories [41, 42], and can be directly used to analyze high dimensional data such as the multi-electrode neural recordings that we studied.

## Acknowledgements

We gratefully acknowledge the support of federal grants NIH BRAIN R01EB026908, NSF NCS-FO 2024364, and of the Swartz Center for Theoretical Neuroscience at the University of Washington. NS is supported by the Klingenstein-Simons Fellows and Pew Biomedical Scholars programs. VB and SB were supported by Simons Foundation MMLS grant 400425. VB was also supported by NIH BRAIN R01EB026945 and NSF Physics Frontiers Center grant PHY 1734030.

